# Bats create a silent frequency band to detect prey through Doppler shift compensation

**DOI:** 10.1101/2025.10.05.680495

**Authors:** Soshi Yoshida, Haruhito Mastumoto, Kohta I. Kobayasi, Shizuko Hiryu

## Abstract

Acquiring information efficiently through sensory inputs is essential for animal survival. Animals with active sensory systems that emit their own signals often optimize the design and use of these signals according to context and purpose^1–6^. In this study, we reveal a previously unrecognized function of Doppler shift compensation (DSC) in bats. During flight, bats actively lower the frequency of their echolocation calls so that echoes remain stable at a reference frequency (*f*_ref_) despite Doppler shifts caused by movement^7–12^. We demonstrate that DSC does not simply serve, as previously thought, to align echoes with the acoustic fovea—a narrow band of maximal auditory sensitivity^13,14^—but in addition, suppresses background noise for detecting prey-derived signals. Using phantom echo playbacks and on-board microphone recordings, we show that bats selectively compensate for the highest-frequency echoes rather than the most intense ones. This process shifts all clutter echoes below *f*_ref_, leaving the spectral band above *f*_ref_ free of stationary-object echoes and secures a “quiet frequency band”. Recordings during prey capture and noise playback experiments revealed that spectral glints from fluttering moth wings appear in this “quiet frequency band” and are exploited for prey detection. This mechanism enhances the high-fidelity detection of prey echoes even in cluttered environments. Such findings reveal a sensory strategy in which animals actively create silence in a critical frequency range. It represents a conceptual advance in active sensing and auditory scene analysis, highlighting how evolution shapes sensory systems to extract reliable information under noisy natural conditions.

## Results and Discussion

### Clarifying the echo selection criteria in DSC behavior

To address the central question of why bats do not compensate for prey echoes, we examined the echo selection rules underlying DSC. In natural flight, bats encounter numerous echoes of varying intensity and Doppler shift, arriving from different directions and distances, and must choose one for compensation. We therefore asked which acoustic feature they prioritize when selecting the target echo. We hypothesized that bats base DSC either on the strongest intensity echo or on the echo with the highest frequency. To test this, we performed a real-time playback experiment. The bats’ calls were captured, processed to manipulate intensity and frequency, and played back as phantom echoes through tweeters (Figure 1A).

**Figure 1.**
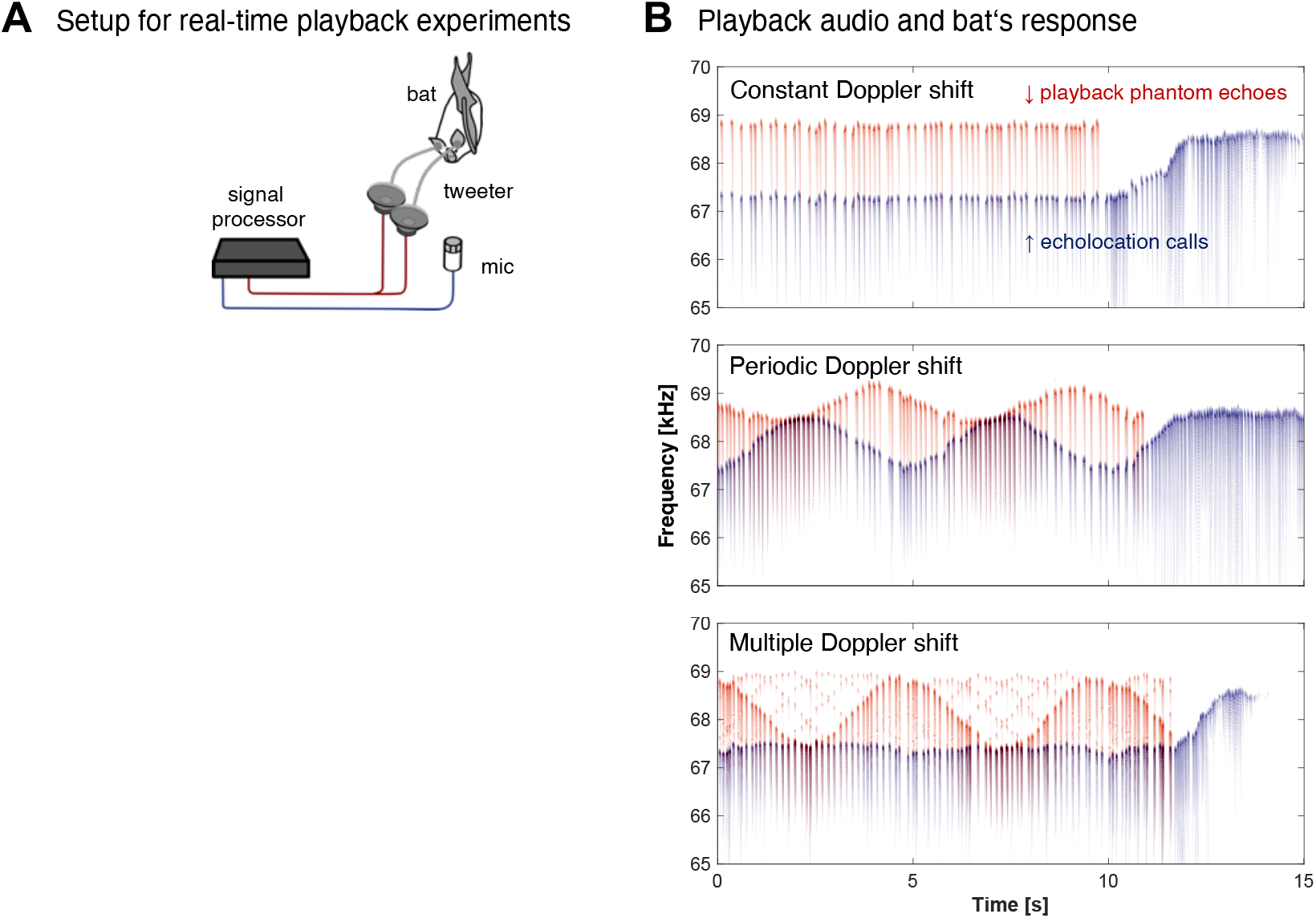
Real-time playback experiment for clarifying the echo selection criteria in DSC behavior. (**A**) Schematic diagram of the experimental setup. The echolocation pulse were captured using a microphone, processed in real-time, and played back through tweeters as phantom echoes (**B**) Representative examples of playback stimuli (red) and bat responses (blue). Under constant Doppler shift conditions, the playback stimulus was processed with a 1.5 kHz frequency shift, and in response, the bats showed a DSC response by consistently lowering their call frequency. Under periodic Doppler shift conditions, where the frequency shift varied sinusoidally between 0 and 1.5 kHz with a 5-second cycle, the bats exhibited a corresponding periodic modulation in their call frequency. Finally, under the multiple Doppler shift condition, where echoes of different intensities were combined such that the maximum-intensity echoes exhibited a frequency shift that modulated periodically while the highest-frequency echoes, though lower in intensity (−20 dB), consistently maintained shift, the bats respond by consistently lowering their call frequency. See also Figure S1.

We first examined two control conditions: a constant Doppler shift and a periodic Doppler shift. Under the constant shift condition, the playback stimulus was processed with a fixed +1.5 kHz frequency shift, and bats consistently lowered their call frequency, demonstrating a clear DSC response. Under the periodic shift condition, where the frequency shift varied sinusoidally between 0 and +1.5 kHz over a 5-s cycle, the bats exhibited a corresponding periodic modulation in their call frequency. Having confirmed that the bats showed stable DSC responses to the control stimuli, we then introduced the multiple Doppler shift condition in four individuals to test how they select target echoes. This stimulus combined echoes that differed in both intensity and frequency shift: the maximum-intensity echoes varied sinusoidally from 0 to +1.5 kHz over a 5 s, while the highest-frequency echoes —20dB weaker— remained fixed around +1.5 kHz. The outcome was clear. Instead of modulating their calls with the sinusoidal shift, bats consistently lowered their call frequency (Figure 1B, see also Figure S1). Thus, bats compensate not for the strongest intensity echoes, but for those with the highest frequency.

### On-board microphones recording of actual echoes perceived by bats

Although phantom echo experiments allow highly flexible stimulus design, they are constrained by artificial conditions. We therefore conducted free-flight experiments using the same four individuals to measure the echoes bats actually receive, in order to validate the plausibility of the findings obtained in the phantom echo experiments. A custom-built on-board microphone was mounted on the bats’ back as they circled in a flight chamber. The chamber had two wall types: “quiet walls,” covered with sound-absorbing material and located outside the nets, and “echoic walls,” left reflective and located in the flight area (Figure 2A). This configuration made the bats to alternately encounter echoic and quiet walls during circular flight, generating a sequence of alternating high- and low-intensity echoes. As in the playback experiments, the maximum-intensity echoes did not necessarily correspond to the highest-frequency ones. Yet the bats consistently compensated for the highest-frequency echoes, maintaining stable call frequencies even when those echoes were weak (Figure 2B, see also Figure S2). The on-board recordings, therefore, confirmed that the DSC strategy observed in real-time playback also occurs under naturalistic flight conditions. Notably, high-quality recordings revealed a distinct spectral structure: the frequencies above the reference frequency *f*_ref_ were strikingly quiet, in contrast to the many prominent echoes below *f*_ref_. This silence appears as a byproduct of compensating for the highest-frequency echoes. The result was a sharply defined “quiet frequency band”, suggesting that bats may use DSC to actively create such a band for prey detection.

**Figure 2.**
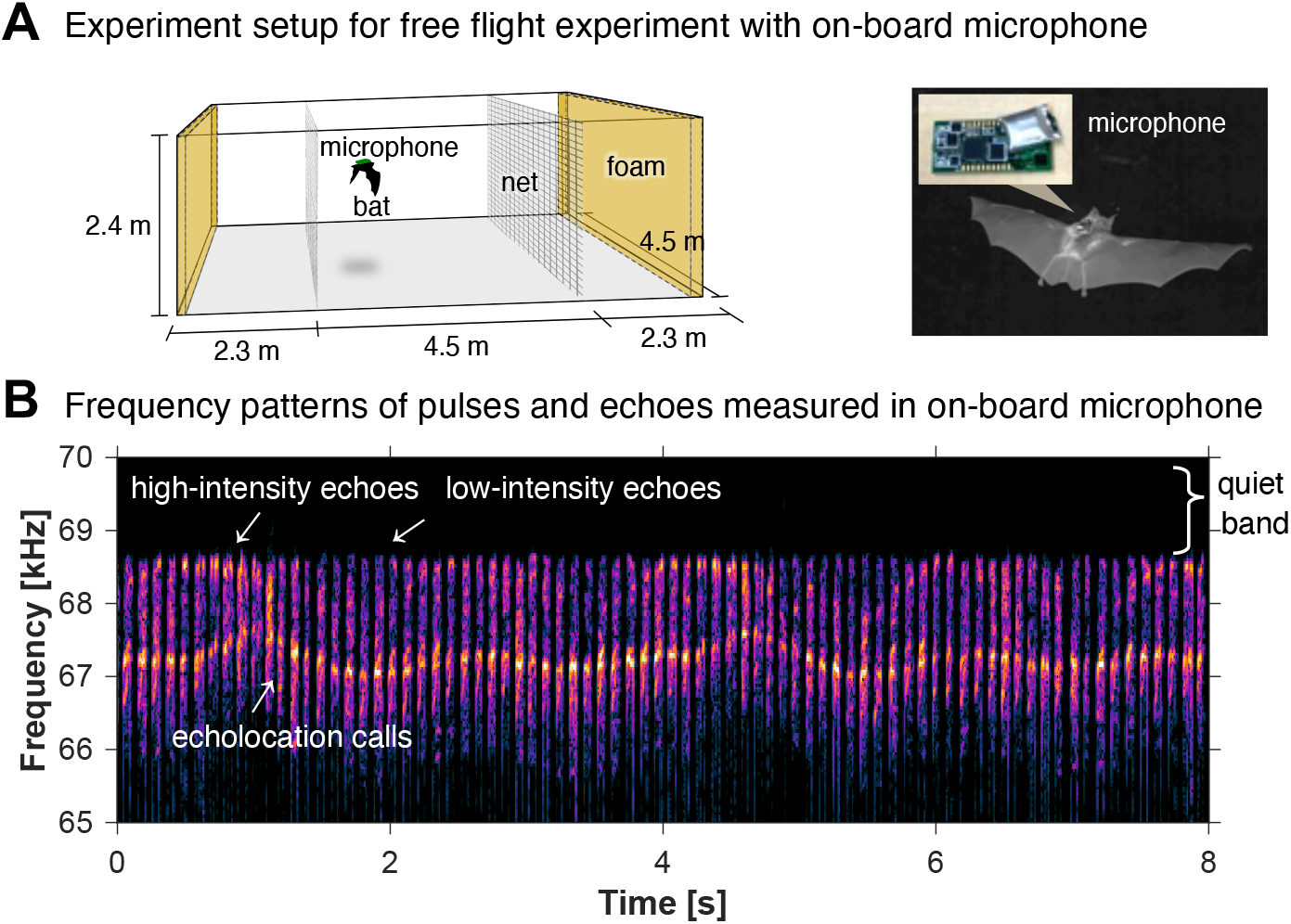
Free flight experiment for echo measurement with on-board microphone. (**A**) Experimental setup and on-board microphone. The flight area was created by nets within a chamber and sound-absorbing material was installed on the walls beyond the nets (quiet walls), while the inner walls remained reflective (echoic walls). This makes bats flying in circles to alternately encounter quiet and echoic wall. (**B**) Representative examples of frequency patterns of pulses and returning echoes recorded by on-board microphones during free flight. A condition was successfully created in which high- and low-intensity echoes alternated, and the maximum-intensity echoes did not necessarily correspond to the highest-frequency ones. The highest-frequency echoes consistently maintained a fixed frequency, confirming that DSC targeted the highest-frequency echoes, not the maximum-intensity echoes, which is consistent with the real-time playback experiment. Furthermore, notably quiet frequency band were observed above the *f*_ref_, whereas numerous echoes were observed below the *f*_ref_. See also Figure S2.

### Measurements of DSC behavior in the context of prey capturing

Next, we measured DSC behavior in the context of prey pursuit. As in the previous experiment, two newly captured wild bats were equipped with the same custom-built on-board microphones, and we recorded the echoes bats received while attacking tethered noctuid moth (Figure 3A). To mimic natural fly-catching, moths were presented to perching bats, which readily took off to attack. Importantly, in the recordings, clear spectral glints appeared in the echoes just before capture (Figure 3B, see also Figure S3). These glints—periodic modulations in amplitude and frequency generated when the CF component of a call reflects from fluttering moth wings— are known to be critical cues for detecting and identifying prey ^15^ (see also Figure S4). Strikingly, the glints occurred with a high signal-to-noise ratio within the “quiet frequency band” identified in the free-flight experiments. These results suggest that DSC does more than simply place echoes into a sensitive band. Instead, it acts as an active strategy to suppress background clutter and carve out a silent spectral channel for prey detection. By stabilizing the frequency of returning echoes, bats effectively create a noise-free spectral window in which prey-generated motion cues are most salient, thereby enhancing the detection of prey-derived spectral glints.

**Figure 3.**
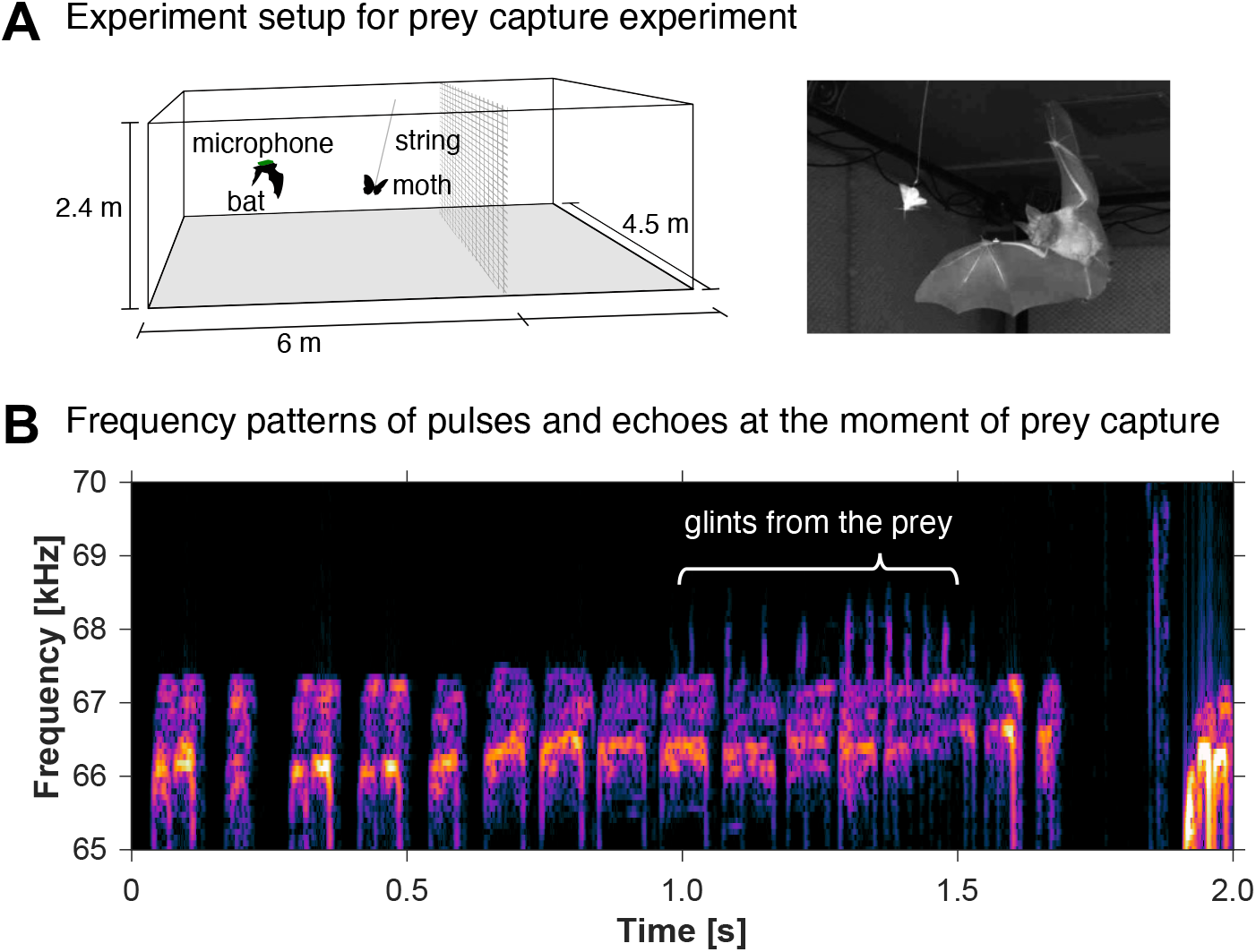
Prey capturing experiment for echo measurement with on-board microphone. (**A**) Experimental setup for prey capturing experiment. A tethered moth was presented, and bats equipped with onboard microphones were released to attack. (**B**) Representative examples of frequency patterns of pulses and returning echoes recorded by on-board microphones at the time of prey capture. Just before the attack, spectral glints caused by the fluttering of moth wings was detected. These glints were observed within the same frequency range as the quiet band identified in the free-flight experiment. See also Figure S3.

### Narrow-band noise playback experiments for prey detection

To further test this hypothesis, we performed a narrow-band noise playback experiment to assess the functional importance of the “quiet frequency band”. Six newly captured wild bats were used in this test. Moths were presented while a narrow band of noise 2 kHz above the resting frequency was played, testing whether bats could still detect and attack prey. As controls, we used a no-noise condition and a low-frequency noise condition in which noise 2 kHz below the resting frequency was presented -this band corresponds to the range where most echoes typically return during flight (Figure 4A).

**Figure 4.**
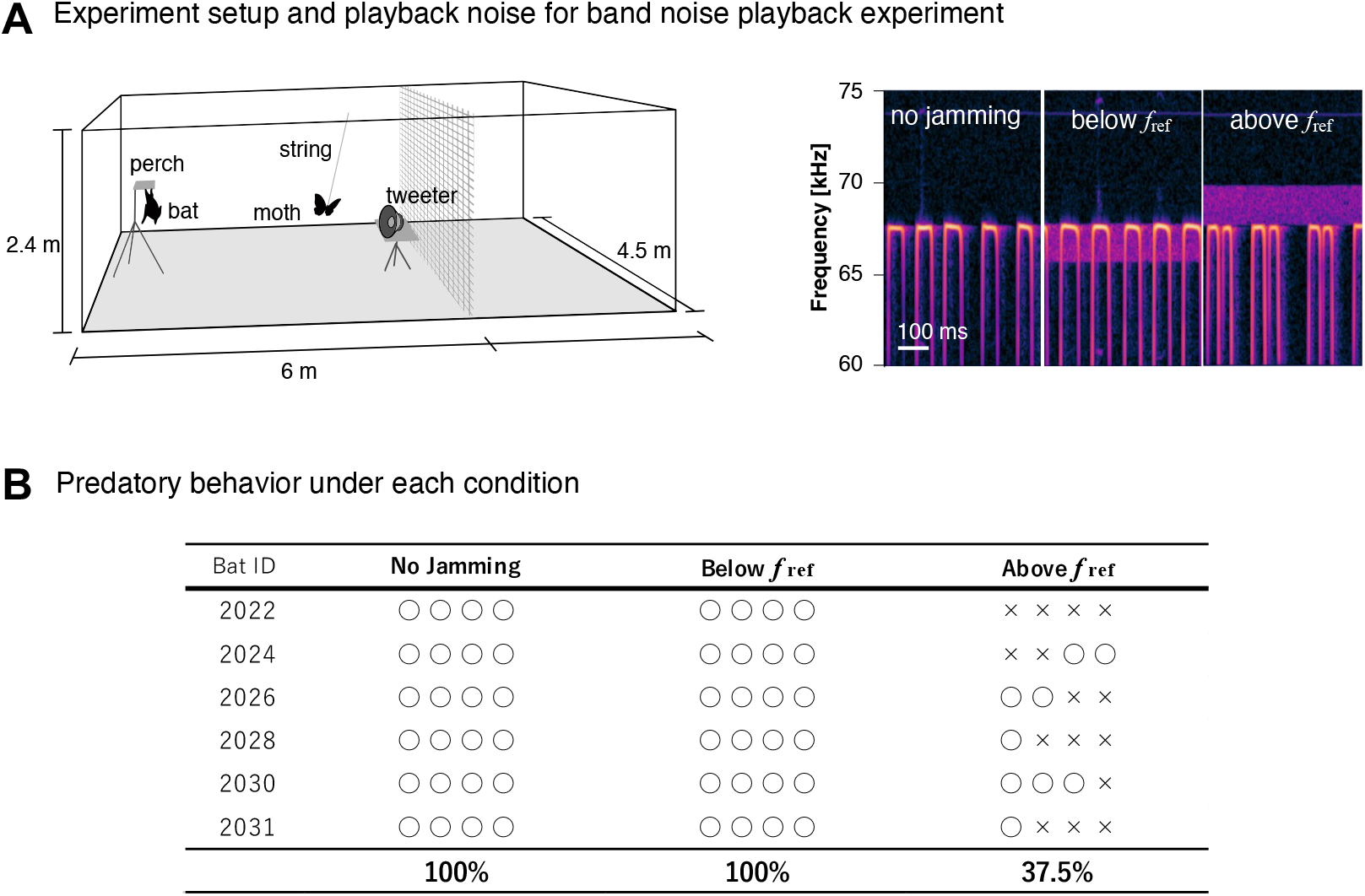
Prey detection test under noise playback in “quiet frequency bands”. Experimental setup for noise playback experiment. The bats were placed on a perch, and moths were presented in front of a tweeter while noise was played back from the tweeter. Three conditions were tested: two controls (no jamming noise and noise 2 kHz below *f*_ref_) and a main condition with noise 2 kHz above *f*_ref_. (**B**) Presence or absence of predatory behavior under each noise condition. Table showing whether bats took off from the perch to capture the moth (○) or remained on the perch (×) after the moth was presented. The order of the four symbols under each condition reflects the actual sequence of presentation. Under control conditions, predatory behavior was observed in 100% of trials. However, when noise was presented at 2 kHz above the *f*_ref_, corresponding to the “quiet frequency band”, the rate dropped to 37.5%. See also Movie S1.

The result was striking. Bats readily attacked under both the no-noise and low-frequency noise conditions, but the attack rate dropped to 37.5% when high-frequency noise jammed the quiet band (Figure 4B, see also Movie S1). These findings demonstrate that bats rely on the quiet frequency band to detect prey-associated spectral glints and that keeping this band free of clutter is essential for successful prey detection.

## Discussion

DSC is one of the most iconic and extensively investigated phenomena in bat research and widely regarded as a hallmark of their remarkable biosonar sophistication ^7–12^. This neural-behavioral specialization provides a powerful model for studying efficient information-acquisition strategies in animals. In active sensing, however, the signal-to-noise ratio (S/N) ultimately constrains performance. Even with finely tuned sensory organs, there are limits to what can be detected. Previous work on DSC during prey pursuit highlighted a paradox: bats compensate not for Doppler shifts in prey echoes, but for those from stationary background objects ^16–18^. If DSC merely served to maximize sensitivity, it should target the echoes of prey— the most relevant signals. Yet, this have not been observed. We therefore hypothesized that DSC serves a broader strategic role. Beyond enhancing auditory sensitivity, bats may exploit the physics of echoes to substantially improve S/N, and we tested this hypothesis experimentally.

Our phantom echo playbacks experiment and on-board microphone recordings show that horseshoe bats compensate for the highest-frequency echoes during DSC, thereby creating a remarkably quiet frequency band above *f*_ref_. Recordings and noise interference in the context of prey capture revealed that spectral glints from fluttering moth wings fall within this quiet frequency band and are exploit for prey detection. These results provide direct evidence for the overlooked function of DSC as S/N improving strategy, showing that HDC bats use DSC not only as a neural specialization but also as a physical strategy to suppress clutter and secure a completely quiet frequency band for prey detection. This finding illustrates how active sensing systems can transcend physiological limits by leveraging physical principles, with implications that extend beyond bats.

Our findings also align with previous reports that auditory sensitivity in HDC bats can extend slightly above *f*_ref_, with relatively shallow decline^19–21^, consistent with the frequency range we identified as the quiet band (Figure S5). Together, these results suggest that the auditory fovea and DSC act in concert: the fovea provides heightened sensitivity around and above *f*_ref_, while DSC eliminates competing echoes from this range.

The discovery that bats combine neural tuning with a physical strategy to create an acoustic “quiet zone” highlights an evolutionary solution to the limits of active sensing. By leveraging physics as well as physiology, bats achieve a functional improvement in signal-to-noise ratio that enhances prey detection under natural clutter. While proximate mechanisms of DSC have been well studied, our work provides a clear explanation of its ultimate function and evolutionary significance, establishing DSC as an adaptive strategy for active sensing in complex environments.

## Acknowledgments

This work was supported by JSPS KAKENHI Grant Number 21H05295 (H.S.), JSPS KAKENHI Grant Number 25H00745 (H.S.), JST SPRING Grant Number JPMJSP2129 (S.Y.), JSPS KAKENHI Grant Number 24KJ2144 (S.Y.)

## Authors’ contributions

**Soshi Yoshida:** Conceptualization: S.Y., H.S. Methodology: S.Y., H.S. Investigation: S.Y., H.M. Visualization: S.Y. Funding acquisition: H.S., S.Y. Project administration: H.S., K.I.K. Supervision: H.S., K.I.K. Writing – original draft: S.Y. Writing – review & editing: S.Y., H.S.

## STAR METHODS

### Key Resource Table

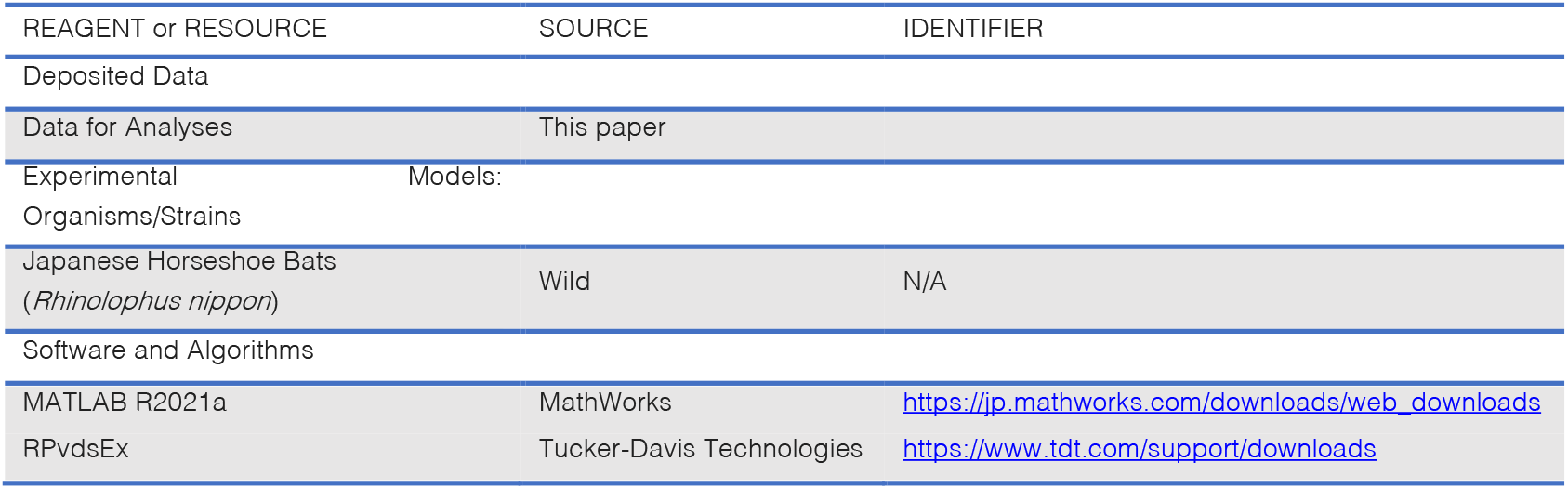

## RESOURCE AVAILABILITY

### Lead contact

Further information and requests for resources should be directed to and will be fulfilled by the lead contact, Soshi Yoshida

### Materials availability

This study did not generate new unique reagents.

### Data and code availability

All data used will be uploaded to Mendeley data

The original code will be uploaded to Mendeley data

Any additional information reported in this work is available from the lead contact upon reasonable request

## EXPERIMENTAL MODEL AND SUBJECT DETAILS

In this study, 11 Japanese horseshoe bats (*Rhinolophus nippon*; 9 males and 2 females) were used. These bats were captured from a cave located in Fukui Prefecture, Japan. We were licensed to collect bats, and we complied with all Japanese laws (permits from Fukui Prefecture in 2022 and 2023). The bats were housed in a room (3 × 2 × 2 m) at Doshisha University in Kyoto, Japan. The temperature and humidity in the room were monitored and controlled to mimic the cave environment, with temperatures around 20 degrees Celsius and humidity above 80% all year round. The room was cleaned, and mealworms and water were placed every 2–3 days to allow the bats to eat and drink *ad libitum*. The experiments were conducted under the permission of the Animal Experiment Committee of Doshisha University (Permission number: A22015, A23017, A24017).

## METHOD DETAILS

### 1. Experiments to elucidate the factors that determine the target echo during DSC

Bats in flight receive multiple echoes with varying intensity and frequency, one of which is selected as the target echo for Doppler shift compensation (DSC). However, it remains unclear what rules bats follow when selecting target echoes from multiple echoes. We hypothesized that bats select either the echo with the maximum-intensity or the one with the highest-frequency as their target among multiple echoes. To test these hypotheses, two experiments were conducted: real-time playback experiment with phantom echoes and a free-flight experiment. Both experiments were conducted in an experimental chamber (9 × 4.5 × 2.4 m). Acoustic foams are put on the walls to minimize echoes from the walls. During the experiments, red LED lighting (wavelengths 650 nm) was used to decrease the possibility of bats using visual information.

#### 1.1 Real-time playback experiments for clarifying the echo selection criteria in DSC behavior

In real-time playback experiments, combinations of multiple phantom echoes with different sound intensities and time-varying frequency shifts were created as stimuli and presented to the stationary bats in real time to test which echoes the bats selected for DSC. Following a protocol of our previous research^22^, real-time playback of phantom echoes was achieved by capturing the bat’s emitted pulses with a ultrasonic microphone (Avisoft, CM16) positioned approximately 20 cm away from the bat (Note that the bats were held by the experimenter to ensure they remained in front of the microphone throughout the experiment). The signals were then sent through a microphone amplifier (TDT, MA3) to a digital signal processor (TDT, RX6, 200 kHz sampling rate) for frequency modulation processing (Doppler shift) using a double heterodyne method. The details of the double heterodyne processing in TDT RX6 correspond to the protocol of our previous study^23^. Since actual echoes, other than the phantom echo signals, are unnecessary as a stimulus for bats, we designed an earphone-like system based on a previous setup^24^, with couplers connected to the tweeter (TDT, ES1) via a speaker driver (TDT, ED1), ensuring that only the playback phantom echoes are heard by the bats (Figure 1A). The frequency response of the playback system was within ±7 dB between 50 and 70 kHz, corresponding to the echolocation call frequency of the Japanese horseshoe bats. The bats’ emitted pulses and playback phantom echoes were recorded using a DAQ (NI, USB-6356, 500 kHz sampling rate). The signal processing delay by TDT RX6 was less than 1 ms, which is sufficiently short.

In this experiment, three different patterns of time-varying frequency shifts were presented for each four bats, processed. The first two patterns served as positive controls, while the last one was the main test stimulus (Figure 1B). The first is a stimulus pattern with a constant +1.5 kHz Doppler shifts from the emitted pulse frequency. The 1.5 kHz Doppler shift corresponds to a typical flight speed of 3-4 m/s with a CF2 frequency of 68 kHz under laboratory conditions^8^. The second stimulus pattern consists of a periodic change in Doppler shifts from 0 to 1.5 kHz with a period of 5 seconds. The third stimulus combined one strong phantom echo with several weak phantom echoes (-20 dB compared to the strong one). This was designed so that if the bat focused on the maximum-intensity echo as the target echo for DSC, the frequency pattern of the stimulus would be perceived as varying sinusoidally from 0 to 1.5 kHz over a 5-second period. In contrast, if the bat focused on the highest-frequency echo for DSC, the stimulus pattern would remain constant at 1.5 kHz without changing over time. Thus, if the bat modulates the CF2 of its emitted pulses sinusoidally over a 5-second period, we can conclude that it is compensating for the echoes with the maximum-intensity. On the other hand, if the bats continuously decrease the frequency of its emitted pulses by the same amount, we can conclude that they are compensating for the highest-frequency echoes.

#### 1.2 Free-flight experiment using on-board microphones to record the actual echoes perceived by bats

For bats in free flight, a more natural behavior, we investigated whether they compensate for the highest-frequency echoes or the maximum-intensity echoes at the reference frequency, *f*_ref_ when performing DSC. A square flight space (4.5 × 4.5 m) was created in the center of the experimental chamber (9 × 4.5 m) by dividing it with two nets (Figure 2A). Sound-absorbing material (attenuation of about -25 dB in the ultrasound range) was installed on the outer walls beyond the nets, creating a contrast in echo intensity between these net-separated walls (‘quiet walls’) and the unobstructed inner walls (‘echoic walls’) within the flight space. In this way, bats flying in circles within the flight space alternated between ‘quiet walls’ and the ‘echoic walls’, creating an environment where the maximum-intensity echoes reaching the bats were not necessarily the highest-frequency echoes. Experiments were conducted on four individual bats.

A custom-made on-board microphone (based on Knowles SPU0410LR5H, logger type, design by Girlie) was attached to the bat’s back to directly measure the echoes from these walls that reach the bats during flight. The weight of the on-board microphone is 2.5 g, including batteries, which is about 10% of the body weight of Japanese horseshoe bats. The sampling rate is 288 kHz, the sensitivity is within ±4 dB between 50 and 70 kHz, and the dynamic range is approximately 70 dB. In particular, within the 65-70 kHz range, which corresponds to the frequency of the CF2 component of the emitted pulse, important for this measurement, the microphone exhibits a ±2.5 dB response and a very low self-noise level equivalent to 20 dB SPL in terms of sound pressure.

### 2. Measurements of DSC behavior during prey capturing

We investigated the DSC behavior performed by bats while they headed towards the flattering prey moths for predation, as well as the echoes reaching them. Furthermore, we examined the contribution of the target-echo determinants identified in Section 1.1 and 1.2 to predatory behavior.

#### 2.1 On-board recordings during tethered moth presentations

The sequence of emitted pulses and echoes received by bats while flying toward a fluttering moth during predation was measured using the same on-board microphone described in Section 1.2 above. The prey moth, *Goniocraspidum pryeri*, was captured in a cave. There are records of bats consuming this moth in the wild^25^. The moth was tethered to the ceiling by a thin wire hooked around its abdomen and a thread (90 cm of #30 cotton yarn, Figure 3A). Without fatally damage, it fluttered vigorously for a few minutes and flew freely within the range of the thread. The tethered moth was gently held by the experimenter and released from their hand after confirming that the bat was stably perched on their perch. The bats equipped with on-board microphone were able to take off from the perches at any time and attack the moths. When a bat successfully caught a moth, it was removed from the wire, allowing the bat to feed on it. Experiments were conducted on five individual bats. Of these, three individuals accepted the on-board microphone, and two successfully attacked tethered moths, providing clear recordings of prey-capture events.

#### 2.2 Narrow-band noise playback experiments for prey detection

Next, to confirm the frequency range crucial for prey detection, narrow-band noise was played at 2 kHz below and above *f*_ref_, respectively. This was done to determine whether the frequency band used by bats for predatory behavior was below or above *f*_ref_. The former is the band where DSC gathers a large number of echoes from the surroundings (Figure 2B, S2), while the latter is the quiet band where no echoes from the surroundings are returned, and only glints from fluttering prey are observed (Figure 3B, S3). The largest spectral glints from *G. pryeri*, are approximately ±1.5 kHz. The playback of 2 kHz band noise completely masks the glints from the moths in the respective bands. Since *f*_ref_ varied across individuals, the resting frequency (*f*_rest_) of each individual was measured the day before the experiment, and band noise was prepared in advance according to each individual’s *f*_rest_. The band noise was output from a DAQ (NI USB-6356, sampling at 500 kHz) and was emitted from a tweeter (Pioneer, PT-R7III) through an audio amplifier (Pioneer, A-D3). Following the same procedure as the previous experiment, the bat was perched on the perch, but the tweeter was positioned 1.5 m behind the moth, directed toward both the bat and the moth (Figure 4A). The sound pressure at the bat on perch was approximately 60 dB SPL. At this perch location, although we could measure the noise from the tweeter, the actual glints from the moth were measured with low sound pressure. Therefore, we assume that this noise is sufficiently louder than the glints. The tethered moth was gently held by the experimenter and released from their hand after confirming that the bat was stably perched on the perch, and after the noise playback had begun. There were three conditions: two controls (one with no noise and one with band noise in the 2 kHz below *f*_ref_) and a main condition with noise in the 2 kHz band above *f*_ref_. Each noise stimulus was pre-generated in MATLAB according to the *f*_ref_ of each individual measured on the previous day. Each condition was repeated in random order four times for each individual. This set of trials was conducted for six individuals in total. We classified cases where the bats took off from the perch toward the moth immediately after its presentation as predatory behavior. Conversely, cases where the bats remained on the perch despite the moth fluttering after being presented were classified as no predatory behavior.

## Reference

1. Donald, G. (1958). Listening in the dark (Yale University Press).

2. Kalko, E.K.V. (1995). Insect pursuit, prey capture and echolocation in pipestirelle bats (Microchiroptera). Anim. Behav. 50, 861–880. 10.1016/0003-3472(95)80090-5.

3. Madsen, P.T., Johnson, M., Aguilar De Soto, N., Zimmer, W.M.X., and Tyack, P. (2005). Biosonar performance of foraging beaked whales (Mesoplodon densirostris). J. Exp. Biol. 208, 181–194. 10.1242/jeb.01327.

4. Hase, K., Kadoya, Y., Maitani, Y., Miyamoto, T., Kobayasi, K.I., and Hiryu, S. (2018). Bats enhance their call identities to solve the cocktail party problem. Commun. Biol. 1, 1–3. 10.1038/s42003-018-0045-3.

5. Gillam, E.H., Ulanovsky, N., and McCracken, G.F. (2007). Rapid jamming avoidance in biosonar. Proc. R. Soc. B Biol. Sci. 274, 651–660. 10.1098/rspb.2006.0047.

6. Bullock, T.H., Behrend, K., and Heiligenberg, W. (1975). Comparison of the jamming avoidance responses in Gymnotoid and Gymnarchid electric fish: A case of convergent evolution of behavior and its sensory basis. J. Comp. Physiol. A 103, 97–121. 10.1007/BF01380047.

7. Schnitzler, H.-U. (1968). Die Uhraschall-Ortungslaute der Hufeisen-Fledermäiuse (Chiroptera-Rhinolophidae) in verschiedenen Orientierungssituationen [The Ultrasonic Sounds of Horseshoe Bats (Chiroptera-Rhinolophidae) in Different Orientation Situations]. Z. Vgl. Physiol. 57, 376–408.

8. Hiryu, S., Shiori, Y., Hosokawa, T., Riquimaroux, H., and Watanabe, Y. (2008). On-board telemetry of emitted sounds from free-flying bats: Compensation for velocity and distance stabilizes echo frequency and amplitude. J. Comp. Physiol. A Neuroethol. Sensory, Neural, Behav. Physiol. 194, 841–851. 10.1007/s00359-008-0355-x.

9. Schoeppler, D., Schnitzler, H.U., and Denzinger, A. (2018). Precise Doppler shift compensation in the hipposiderid bat, Hipposideros armiger. Sci. Rep. 8, 1–11. 10.1038/s41598-018-22880-y.

10. Long, G.R., and Schnitzler, H.U. (1975). Behavioural audiograms from the bat, Rhinolophus ferrumequinum. J. Comp. Physiol. A 100, 211–219. 10.1007/BF00614531.

11. Gaioni, S.J., Riquimaroux, H., and Suga, N. (1990). Biosonar behavior of mustached bats swung on a pendulum prior to cortical ablation. J. Neurophysiol. 64, 1801–1817. 10.1152/jn.1990.64.6.1801.

12. Schnitzler, H.U., and Denzinger, A. (2011). Auditory fovea and Doppler shift compensation: Adaptations for flutter detection in echolocating bats using CF-FM signals. J. Comp. Physiol. A Neuroethol. Sensory, Neural, Behav. Physiol. 197, 541–559. 10.1007/s00359-010-0569-6.

13. Schuller, G., and Pollak, G. (1979). Disproportionate frequency representation in the inferior colliculus of doppler-compensating Greater Horseshoe bats: Evidence for an acoustic fovea. J. Comp. Physiol. A 132, 47–54. 10.1007/BF00617731.

14. Schuller, G. (1984). Natural ultrasonic echoes from wing beating insects are encoded by collicular neurons in the CF-FM bat, Rhinolophus ferrumequinum. J. Comp. Physiol. A 155, 121–128. 10.1007/BF00610937.

15. von der Emde, G., and Menne, D. (1989). Discrimination of insect wingbeat-frequencies by the bat Rhinolophus ferrumequinum. J. Comp. Physiol. A 164, 663–671. 10.1007/BF00614509.

16. Schnitzler, H.U., and Flieger, E. (1983). Detection of oscillating target movements by echolocation in the Greater Horseshoe bat. J. Comp. Physiol. A 153, 385–391. 10.1007/BF00612592.

17. Mantani, S., Hiryu, S., Fujioka, E., Matsuta, N., Riquimaroux, H., and Watanabe, Y. (2012). Echolocation behavior of the Japanese horseshoe bat in pursuit of fluttering prey. J. Comp. Physiol. A Neuroethol. Sensory, Neural, Behav. Physiol. 198, 741–751. 10.1007/s00359-012-0744-z.

18. Ma, N., Xia, H., Yu, C., Wei, T., Yin, K., and Luo, J. (2024). Effects of insect pursuit on the Doppler shift compensation in a hipposiderid bat. J. Exp. Biol. 227. 10.1242/jeb.246355.

19. Taniguchi, I. (1985). Echolocation sounds and hearing of the greater Japanese horseshoe bat (Rhinolophus ferrumequinum nippon). J. Comp. Physiol. A 156, 185–188. 10.1007/BF00610860.

20. Neuweiler, G. (1970). Neurophysiologische Untersuchungen zum Echoortungssystem der Großen Hufeisennase Rhinolophus ferrum equinum Schreber, 1774. Z. Vgl. Physiol. 67, 273–306. 10.1007/BF00340953.

21. Schoeppler, D., Kost, K., Schnitzler, H.U., and Denzinger, A. (2023). Transmitter and receiver of the low frequency horseshoe bat Rhinolophus paradoxolophus are functionally matched for fluttering target detection. J. Comp. Physiol. A Neuroethol. Sensory, Neural, Behav. Physiol. 209, 191–202. 10.1007/s00359-022-01571-0.

22. Yoshida, S., Hase, K., Kobayasi, K.I., and Hiryu, S. (2024). Protocol for the playback of phantom echoes from approaching objects to scanning bats. STAR Protoc. 5, 103453. 10.1016/j.xpro.2024.103453.

23. Yoshida, S., Hase, K., Heim, O., Kobayasi, K.I., and Hiryu, S. (2024). Doppler detection triggers instantaneous escape behavior in scanning bats. iScience 27, 109222. 10.1016/j.isci.2024.109222.

24. Schuller, G. (1997). A cheap earphone for small animals with good frequency response in the ultrasonic frequency range. J. Neurosci. Methods 71, 187–190. 10.1016/S0165-0270(96)00142-2.

25. Sano, A. (2006). Impact of predation by a cave-dwelling bat, Rhinolophus ferrumequinum, on the diapausing population of a troglophilic moth, Goniocraspidum preyeri. Ecol. Res. 21, 321–324. 10.1007/s11284-005-0122-1.

