## Supplementary PDF for "Bats create a silent frequency band to detect prey through Doppler shift compensation"

**This PDF file includes:**

Supplementary Results and Text

Figures S1 and S5

Captions for Movies S1

**Other Supplementary Materials for this manuscript include the following:**

Movies S1

### Supplementary Results and Text

The present study indicate that DSC creates a “quiet frequency range” that may be utilized for detecting glints from the prey. To further confirm this finding, we conducted two additional experiment to verify that bats primarily rely on spectral glints, rather than amplitude glints, for prey detection, and that these spectral glints are detected within the quiet frequency band generated by DSC.

#### Verification of the importance of spectral glint in prey detection

Although both types of glints physically occur in echoes from fluttering prey, establishing the dominant role of spectral glints is critical for supporting our hypothesis. We presented phantom echoes convolved with each of three different stimuli; (1) frequency and amplitude glints generated by the fluttering moths, (2) artificially created frequency-only glints, and (3) artificially created amplitude-only glints. For stimulus (1), actual glints were measured by transmitting a pure tone at 68 kHz corresponding to the typical frequency of the CF2 component in the Japanese horseshoe bat, to a fluttering moth, using a tweeter (Pioneer, PT-R7III), an audio amplifier (Pioneer, A-D3), a microphone (Titley Scientific, Anabat SD2) and a DAQ (NI USB-6356, sampling at 500 kHz). The moth was affixed to a thin rod with beeswax and positioned approximately 10 cm from the tweeter and the microphone which were placed side by side close alignment. Representative glints were then extracted from recorded echoes, which clearly included both frequency and amplitude glints. For stimuli (2), amplitude glints with the same modulation depth and interval (10 dB, 35 ms), mimicking the amplitude component of the measured glints, and for stimuli (3), spectral glints with the same modulation width and interval (1.5 kHz, 35 ms), mimicking the frequency component, were generated. These were then repeatedly embedded into the emitted pulse from a perching bat using the double heterodyne technique and played back in real time as phantom echoes from the moths. The tweeter and microphone were placed side by side and positioned 1.5 m from the bat's perch same as in the previous predation experiments described in section 3 (Figure S4A, B). For the five individuals that had exhibited predatory behavior toward real moths just before this experiment, we conducted these three types of stimuli four times each, totaling 12 trials per individual. Since the moths were not physically present, complete predatory behavior could not be observed. Thus, we recorded the take-off from the perch immediately following the glint presentation as an instance of predatory behavior. As a result, bats exhibited attacks in 85% of trials with (1) measured glints, 75% with (2) frequency-only glints, and 0% with (3) amplitude-only glints (Figure S4C). These results demonstrate that bats primarily rely on spectral glints for prey detection.

Measurement of auditory sensitivity around the $f_{ref}$

The auditory sensitivity of bats was measured. Specifically, we focused on frequencies around $f_{ref}$, to assess detailed auditory sensitivity through behavioral auditory response analysis. Based on previous studies, the audiogram was measured using the Pleyel reflex behavior, in which bats flick their pinnae when they hear a sound [19,21]. Bats were placed in a soundproof box (50 × 50 × 100 cm) and kept at rest until they stopped emitting pulses and moving their pinnae. The perch inside the soundproof box was placed 50 cm away from the tweeter on the floor and was intentionally made very small (2 cm) to ensure that the bat’s relative position to the tweeter remained largely unchanged, even if the bat attempted to move. Bat behavior was monitored using an IR camera (DMK 33UX273, frame rate at 30fps) synchronized with the sound presentation system. Tone bursts of 30 ms duration with a 1.5 ms rise and fall time were generated with at various frequencies (1kHz step between 63-66 kHz and 71-73 kHz, 0.5 kHz step between 66-71 kHz) and sound pressures levels (1 dB step). The tones were presented through a tweeter (TDT, ES1), via a DAQ (NI USB-6356, sampling at 500 kHz), and amplified (TDT, ED1). The bats' response, specifically their pinna movements immediately after the tone bursts were tracked using DeepLabCut, a marker less tracking software for animals (Figure S5A). This experiment was conducted on three individual bats. The results indicate that auditory sensitivity is maintained at levels comparable to that at $f_{\mathrm{ref}}$, and in some cases may even appear slightly enhanced, within the range extending 2–3 kHz above $f_{\mathrm{ref}}$ (Figure S5B).

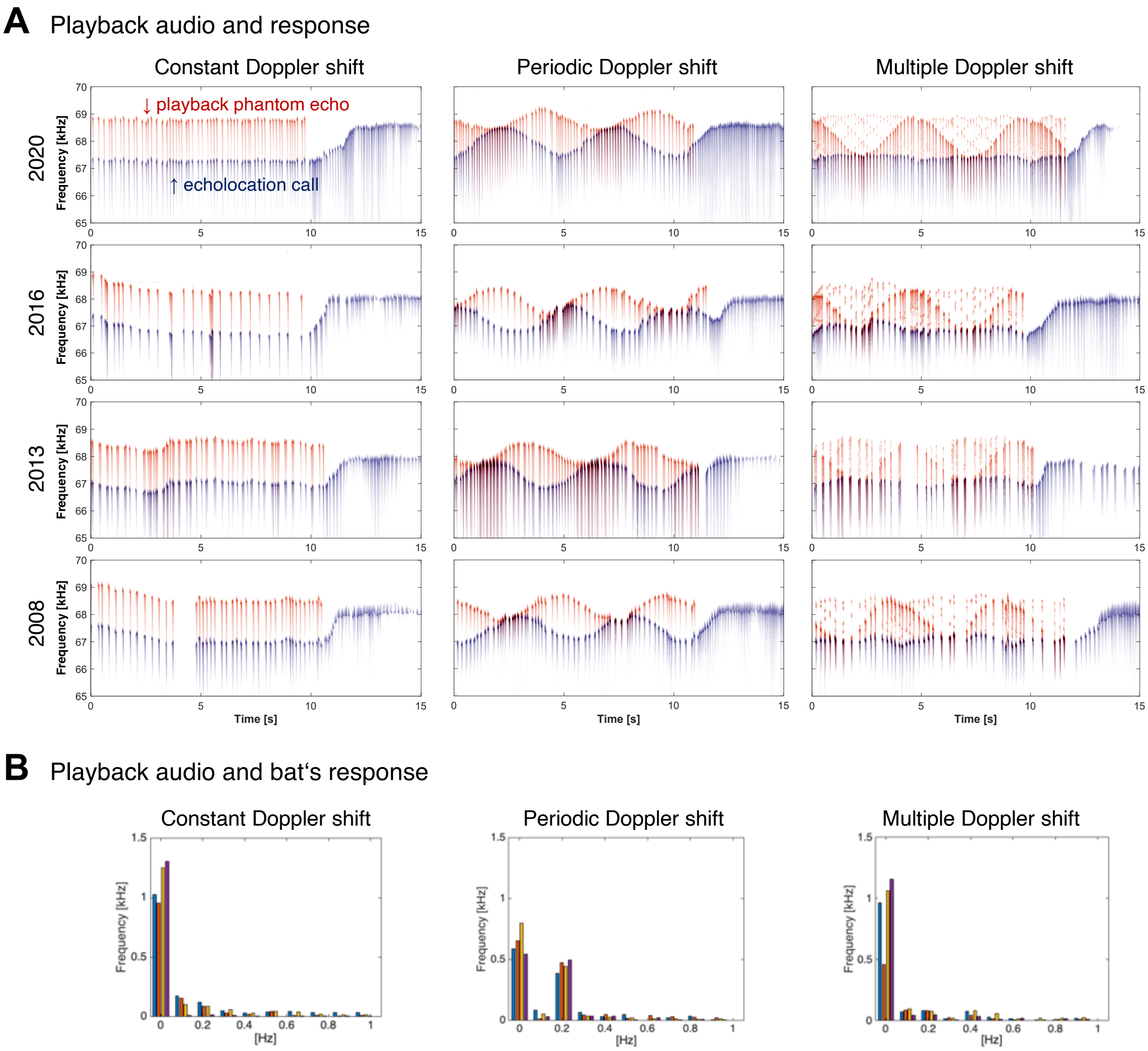


**Figure S1. Group data from all four bats in the real-time playback experiment.**

(**A**) Playback stimuli (red) and bat responses (blue) in all four individuals. Under both constant and multiple Doppler shift conditions, all bats showed a DSC response by consistently lowering their call frequency. Under periodic Doppler shift conditions, they exhibited a DSC response that varied sinusoidally with a 5-second cycle. (**B**) FFT spectra of 10-s time series of frequency differences between call frequency and $f_{ref}$, obtained to quantify periodic changes in call frequency. This analysis allows us to detect both steady offsets (0 Hz) and 5-s periodic components (0.2 Hz). Under constant and multiple Doppler shift conditions, only the 0 Hz peak was observed, while under the periodic condition, an additional 0.2 Hz peak was evident, indicating a 5-s periodic modulation.

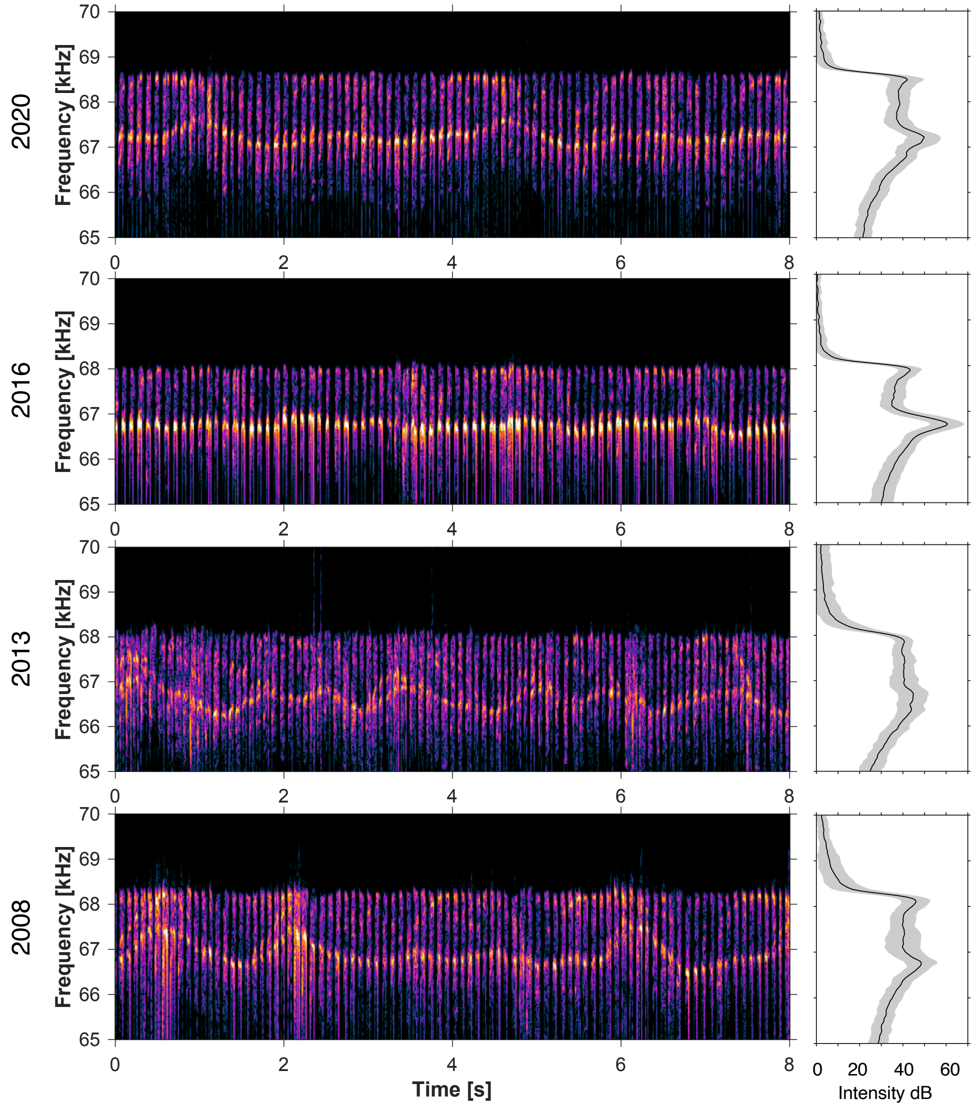


**Figure S2. Group data of free flight recordings with on-board microphones.** Frequency patterns of pulses and returning echoes recorded from all four individuals. The highest-frequency echoes consistently maintained a fixed frequency, consistent with the real-time playback experiment. The spectrum represents intensity with reference to the microphone noise level (about 20 dB SPL). Importantly, a notably quiet frequency band above $f_{ref}$ was clearly observed in all individuals, where the signal intensity approached the microphone noise level, confirming that DSC consistently produced this exceptionally silent region across subjects.

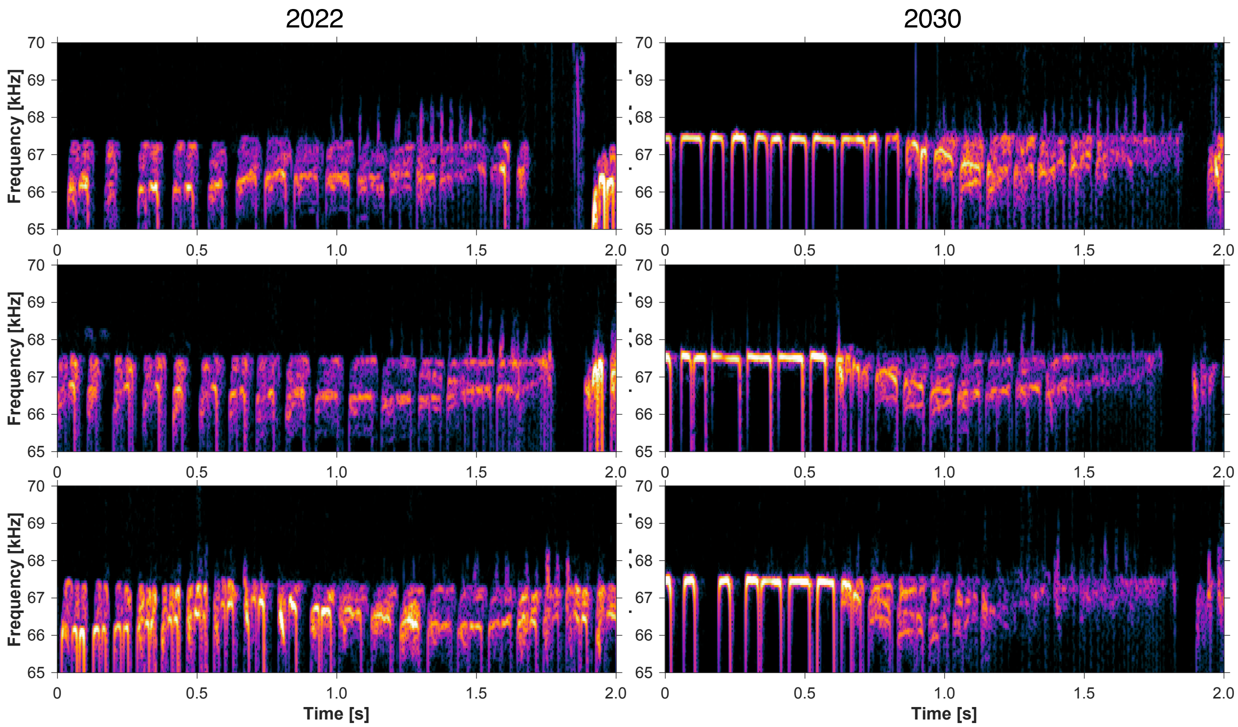


**Figure S3. Spectrograms of pulses and echoes during prey capture flights.**

Frequency patterns of pulses and returning echoes from two bats across three prey capture scenes (six in total). In all cases, spectral glints generated by moth wing flutter were clearly observed immediately before the attack. Importantly, these glints consistently appeared within the same frequency range as the quiet band identified in the free-flight experiment, confirming that this spectral feature was robust across individuals and trials.

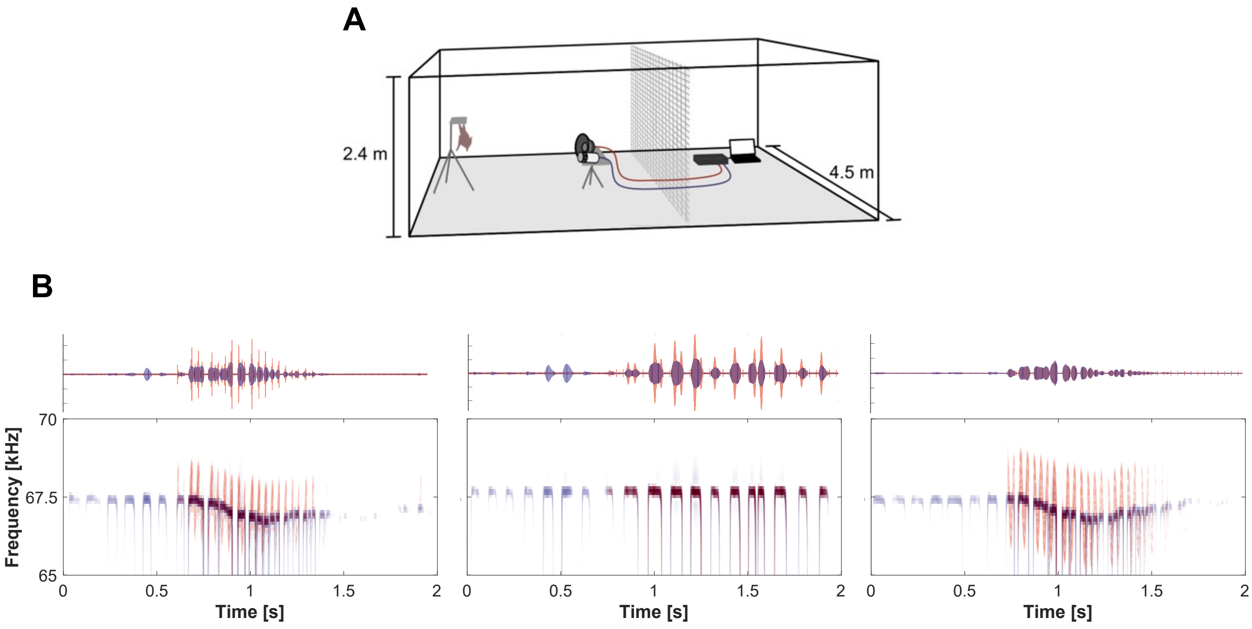


**C**

| Bat ID | Real moth | Recorded glint | Amp. glint | Freq. glint |
|---|---|---|---|---|
| 2022 | ○ | ○○○× | ×××× | ○○○× |
| 2024 | ○ | ○○×○ | ×××× | ○○×× |
| 2026 | ○ | ○○○○ | ×××× | ○×○○ |
| 2028 | ○ | ○○○× | ×××× | ○○○× |
| 2030 | ○ | ○○○○ | ×××× | ○○○○ |
| 2031 | × | - | - | - |
| | 83% | 85% | 0% | 75% |

**Figure S4. Verification of the importance of spectral glint in prey detection.**

(**A**) Experimental setup for real-time playback experiment. Emitted calls captured by microphones are processed and played back to bats in real time. (**B**) Three types of playback stimuli (red): convolution with actual glints measured from moths, addition of amplitude glints only, and addition of spectral glints only. (**C**) The bats flew from the perch toward the tweeter in response to true glint and frequency-only glint but showed no reaction to amplitude-only glint.

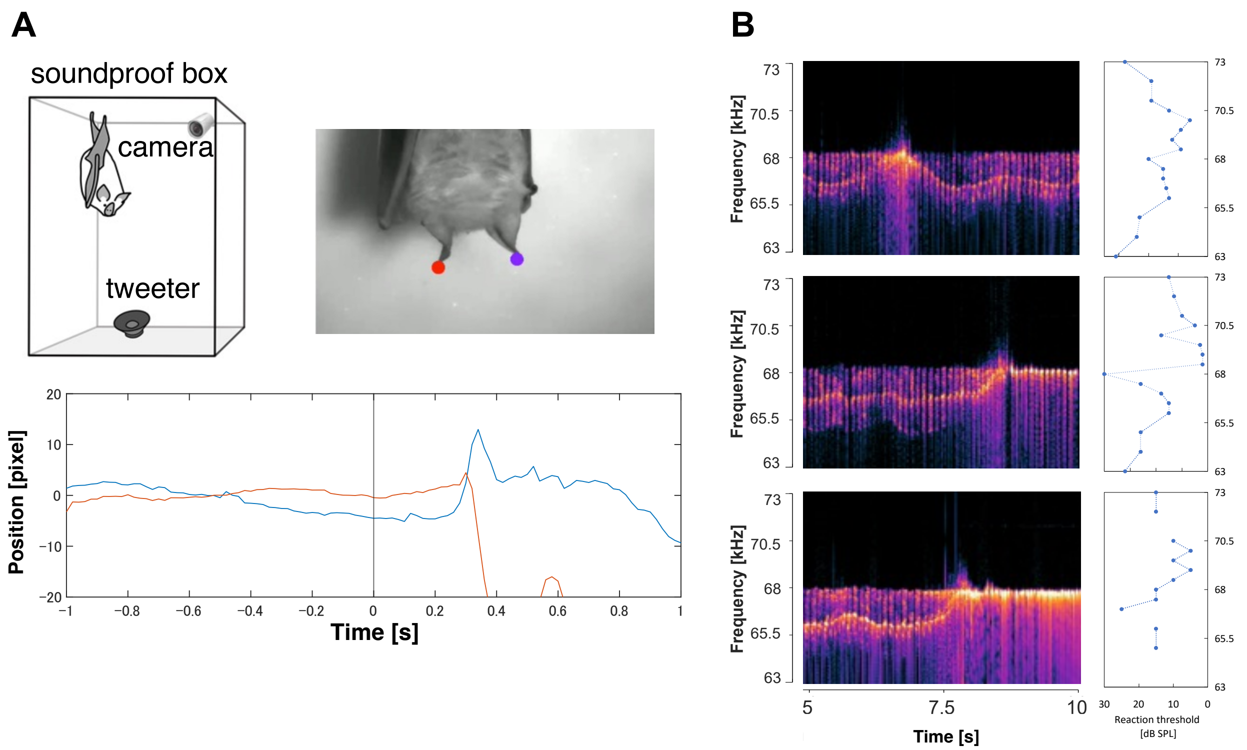


**Figure S5. Measurement of auditory sensitivity around the $f_{ref}$.**

(A) Experimental setup for measuring auditory sensitivity. Sound stimuli were presented from a tweeter to bats sitting on a perch inside a soundproof box. Pinna coordinates were extracted from video recordings to assess whether the ears moved immediately after stimulus presentation. The figure below illustrates a typical example of ear coordinate traces when a reaction occurs. (B) Auditory sensitivity curves obtained by examining the pinna response under various sound pressures and frequencies for three individuals. Auditory sensitivity is high around $f_{ref}$, and there is a tendency for sensitivity to be slightly better in the higher frequency range, where there are no echoes, than in the slightly lower frequency range, where there are many echoes, relative to $f_{ref}$. The results indicate that auditory sensitivity is maintained at levels comparable to that at $f_{ref}$, and in some cases may even appear slightly enhanced, within the range extending 2–3 kHz above $f_{ref}$.

**Movie S1. Prey detection test under noise playback in "quiet frequency bands".** Example of Bat 2028's prey-attack behavior under the three noise presentation conditions. The video shows a 3-second period immediately after moth presentation, replayed at one-quarter speed (final duration: 12 seconds).
